# Direction-aware functional class scoring enrichment analysis of Infinium DNA methylation data

**DOI:** 10.1101/2024.02.22.581670

**Authors:** Mark Ziemann, Mandhri Abeysooriya, Anusuiya Bora, Séverine Lamon, Mary Sravya Kasu, Mitchell W. Norris, Yen Ting Wong, Jeffrey M. Craig

**Affiliations:** Burnet Institute, Melbourne, Australia; Deakin University, School of Life and Environmental Sciences, Geelong, Australia; Deakin University, School of Exercise and Nutrition Sciences, Institute for Physical Activity and Nutrition, Geelong, Australia; Deakin University, School of Medicine, Geelong, Australia; Murdoch Children’s Research Institute, Melbourne, Australia

**Keywords:** Pathway analysis, functional enrichment analysis, Infinium Array, DNA methylation, epigenetics, epigenome-wide association study

## Abstract

Infinium Methylation BeadChip arrays remain one of the most popular platforms for epigenome-wide association studies, but tools for downstream pathway analysis have their limitations. Functional class scoring (FCS) is a group of pathway enrichment techniques that involve the ranking of genes and evaluation of their collective regulation in biological systems, but the implementations described for Infinium methylation array data do not retain direction information, which is important for mechanistic understanding of genomic regulation. Here, we evaluate several candidate FCS methods that retain directional information. According to simulation results, the best-performing method involves the mean aggregation of probe limma t-statistics by gene followed by a rank-ANOVA enrichment test using the mitch package. This method, which we call “LAM”, outperformed an existing over-representation analysis method in simulations, and showed higher sensitivity and robustness in an analysis of real lung tumour-normal paired datasets. Using matched RNA-seq data we examine the relationship of methylation differences at promoters and gene bodies with RNA expression at the level of pathways in lung cancer. To demonstrate the utility of our approach, we apply it to three other contexts where public data were available. Firstly, we examine differential pathway methylation associated with chronological age. Secondly, we investigate pathway methylation differences in infants conceived with in vitro fertilisation. Lastly, we analyse differential pathway methylation in 19 disease states, identifying hundreds of novel associations. These results show LAM is a powerful method for the detection of differential pathway methylation as compared to existing methods. A reproducible vignette is provided to illustrate how to implement this method.

## Introduction

DNA methylation is the most widely studied epigenetic mark, for its role in development and disease.^1^ Hundreds of epigenome-wide association studies (EWASs) are conducted each year to understand DNA methylation patterns in disease and other contexts.^2^ Infinium arrays remain the preferred platform for EWASs due to low cost and analytical simplicity as compared to high throughput methylation sequencing.^3^ Infinium arrays typically include multiple probes per gene in different locations including promoters, CpG islands, gene bodies and enhancers,^4^ which complicates downstream functional enrichment analysis, otherwise known as pathway analysis or gene set enrichment analysis. Enrichment analysis comes in two popular types: over-representation analysis (ORA) and functional class scoring (FCS).^5^ A third type of enrichment analysis called pathway topology improves upon other methods with more sophisticated modelling of gene network activations, but these are yet to be adopted widely.^6^ In ORA, genes with probes that meet an arbitrary significance threshold are selected and compared to a background list of all genes measured in the assay. The test seeks to identify sets of genes (e.g., ontologies, pathways) that are over-represented in the gene list of interest relative to the background.^7^ FCS takes a different approach by ranking all detected genes by a differential regulation score (e.g., fold change, confident effect size, t-statistic) followed by a test to assess whether each set of genes has a distribution of scores that is different from the null.^8^ The gsameth() function of the missMethyl package is the state-of-the-art method for ORA of Infinium methylation array data as it addresses issues related to probes belonging to more than one gene and the fact that one gene can have multiple probes.^9^ As gsameth is an ORA method, results strongly depend on the significance threshold used,^10^ and also on the proportion of probes that meet this threshold. FCS methods ebGSEA and methylGSA have been developed for EWAS data and are suggested to have better sensitivity to determine subtle associations between pathways and differential methylation.^10,11^ The ebGSEA tool uses an empirical Bayes modeling approach followed by a one-tailed nonparametric test for gene set enrichment.^11^ The methylGSA package implements two different approaches. Firstly, the methylRRA function, which uses robust rank aggregation for detecting differentially methylated genes taking into consideration the variable number of probes followed by an FCS test based on z-scores.^10^ Secondly, methylglm uses a logistic regression approach to model differential methylation for genes inside and outside each gene set.^10^ These methods are indeed more sensitive than ORA approaches, but as they are based on one-tailed tests irrespective of the direction of methylation change, it limits the utility for downstream interpretation of the importance of methylation changes on genomic regulation. Moreover, as pathways tend to be either upregulated or downregulated in omics assays, combining the two tends to dilute their signal and reduce sensitivity.^12^ Neither gsameth nor methylGSA report computed enrichment scores, which hinders downstream interpretation, as the enrichment score is a useful surrogate for effect size. Without an enrichment score, users are limited to using statistical significance values to prioritise results, which biases towards large gene sets that have non-specific biological functions.

Here we aim to develop and evaluate methods for two-tailed FCS of Infinium methylation array data that addresses these limitations. Since genes have a variable number of probes, there are many conceivable ways that FCS could be applied to Infinium methylation array data. The first set of methods (type I) uses differential probe methylation data from limma^13^ as an input. The differential methylation values of probes belonging to each gene are scored and these gene scores undergo an enrichment test. The second set of methods (type II) first aggregate probe methylation values for each gene before differential methylation analysis with limma. Then, differential gene methylation results are used for a downstream enrichment test. With the above methods, there are several methodological choices, such as the type of aggregation, and the type of downstream test. To determine the best approach to applying FCS to Infinium methylation data, in this study, we use simulated data and compare results to existing ORA methods. We then examine the sensitivity of selected methods on real cancer data, investigate the association with gene expression and end with additional examples related to human aging, assisted reproductive technologies and a large-scale EWAS of 19 diseases.

## Methods

### Implementation overview

Functional enrichment analysis is a process of data summarisation from genes to gene sets (pathways). This is made more complicated for Infinium methylation array data due to the presence of multiple probes per gene, meaning the data needs to be summarised from probes to genes and then to gene sets. Here, we outline eight potential approaches for FCS of Infinium methylation data implemented in R (v4.3.2), where methods 1-5 are type I and 6-8 are type II:

1. Limma Average t-test (LAT). Differential methylation analysis is conducted at the probe level with limma (v3.58.1), and the limma t-statistics for each gene are summarised (arithmetic mean). The mean t-statistics are used in a downstream two-sample two-way t-test of gene set enrichment.
2. Limma Top t-test (LTT). As above, except instead of calculating the mean t-statistic, the probe with the largest magnitude is selected to represent the gene.
3. Limma Average Wilcox (LAW). Similar to LAT, except instead of the two-sided t-test of gene set enrichment, a non-parametric alternative, the Wilcoxon signed-rank test is used.
4. Limma Average Mitch (LAM). Similar to LAW, however, instead of the Wilcoxon test, the mitch package^14^ (v1.15.0) is used to execute a two-way, two-sample ANOVA-on-ranks test as described previously^15^.
5. Limma Rank Mitch (LRM). Probes are ranked by t-statistic. The mean rank values for probes belonging to each gene are computed and used as input for a two-way, two-sample ANOVA-on-ranks test.
6. Aggregate Limma t-test (ALT). In this method, all probe measurements belonging to each gene are averaged before conducting differential methylation analysis with limma. The limma t-statistics are then used downstream for enrichment analysis with the two-sample two-way t-test.
7. Aggregate Limma Wilcox (ALW). Similar to ALT, however, it uses the Wilcoxon non-parametric test for gene set enrichment.
8. Aggregate Limma Mitch (ALM). Similar to ALW, however, it uses the Mitch package to execute a two-way, two-sample ANOVA-on-ranks test.

The above were compared to a standard ORA-based approach (GSA), which involves limma on probes, selection of statistically significant probes (FDR<0.05) for ORA with the gsameth() function of missMethyl (v1.36.0) which conducts a modified hypergeometric test that accounts for multiple probe biases. In the case that fewer than 250 significant probes were identified, the 250 probes with the smallest p-values were selected. We conducted separate tests for increased and decreased probes, and we specified the background as all probes that passed quality control filtering.

### Method validation using simulations

To assess these methods, we adopted a simulated data approach based on the selected modification of real methylation data. We downloaded raw intensity EPIC IDAT files from NCBI Gene Expression Omnibus (GEO) for study GSE158422, which consists of lung tumour and normal adjacent tissues from 37 patients.^16^ For the simulations, only non-cancerous datasets were used. Probe annotations were obtained from the “IlluminaHumanMethylationEPICanno.ilm10b4.hg19” Bioconductor package. We randomly sampled datasets to serve as control and cases (no replacement), with group sizes varying between 3 and 12. One thousand random gene sets were created with sizes varying between 20 and 100 members, with member genes drawn from the annotation set. Random gene sets were used to avoid problems caused by the large overlap among real gene sets. Throughout our evaluations, 50 gene sets were selected to be differentially methylated, 25 with increased and 25 with decreased methylation. From these gene sets, half of the member genes were selected. For those selected genes, half of the annotated probes were selected. We adjusted the M-values in the case group by a specified amount, which we call the “delta”, which we varied between 0.1 and 0.5. Following the incorporation of methylation changes to selected probes in the case group, the data underwent limma differential methylation analysis, followed by enrichment analysis using gsameth, or one of the eight FCS methods outlined above. Enrichment results were then filtered for FDR<0.05 and the correct direction of methylation change. Differentially methylated gene sets observed were compared to gene sets selected to be the ground truth so that the number of true positives (TP), false positives (FP) and false negatives (FN) could be determined. At each setting for group size, delta and gene set size, 100 replications were conducted, each with a set seed to ensure reproducibility. Simulations were conducted on a SLURM high-performance computing cluster.^17^ Mean TP, FP and FN values were determined, and then precision, recall and F1 scores were calculated.

### Sensitivity analysis

The full GSE158422 dataset including normal and cancer samples of all 37 patients underwent differential analysis with limma correcting for patient-of-origin effects. Downstream pathway enrichment analysis was conducted using the GSA and LAM methods with Reactome pathways downloaded from MSigDB website (version 2023.1)^18^. LAM relies on probe-gene associations from the “UCSC_RefGene_Name” column of the “IlluminaHumanMethylationEPICanno.ilm10b4.hg19” annotation. As gene annotations are several years old, we used HGNChelper to update them (database current as of 21-Dec-2023)^19^ (3,253 gene symbols were updated). Euler diagrams were created with the eulerr R package (v7.0.0). To test sensitivity, a random subset of n patients was selected followed by pathway enrichment analysis with the respective method, with n varying between 2 and 30. Significantly enriched pathways were defined as those with FDR values < 0.05. The subset significant results were compared to the full group (n=37) results to test the sensitivity of these enrichment methods, and whether findings at a smaller sample size were consistent with the full group. This process was repeated 50 times for each sample size.

### Association of methylation with gene expression

RNA-seq gene counts for the same set of lung cancer patients were downloaded from GEO (accession: GSE158420). HGNChelper was used to fix gene names converted to dates, followed by DESeq2 differential analysis. This underwent filtering to remove genes with expression below 10 reads per sample on average across the dataset. Then DESeq2^20^ was used to compare normal and cancerous tissue gene expression taking into consideration sample pairing. The differential expression results underwent enrichment analysis using the mitch package^14^ with default settings for DESeq2 data tables. CpG sites annotated as promoters were considered separately from those located at gene bodies in mitch analysis. For comparison, GSA was used to separately examine pathways using separate analyses of promoter and gene body CpGs. For this analysis, Reactome gene sets were used. Subsequently, to generate the pathway heatmap, we used mitch in multivariate mode by analysing differential RNA expression, promoter and gene body methylation together, prioritising the results by S distance, a surrogate measure of effect size.

### Pathway-level differential methylation in aging

In order to demonstrate the utility of the LAM method for large EWAS studies, we examined EPIC methylation profile associations with age in two independent cohorts involving a total of 7,036 participants.^21^ Limma summary statistics for chronological age including discovery and replication groups were read into R and enrichment analysis was conducted with the LAM method. Briefly, gene-probe associations were first obtained from the “IlluminaHumanMethylationEPICanno.ilm10b4.hg19” dataset, followed by an update of gene symbols with HGNChelper. Discovery and replication data were imported separately with mitch and multivariate enrichment analysis was conducted with gene sets from Reactome.

### Pathway-level differential methylation in assisted reproductive technology

In order to demonstrate that methylation patterns across different array platforms can be compared using the LAM method, we examined methylation differences in infants associated with assisted reproduction as described by two independent studies. The Estill study was published in 2016 and is based on the HM450K array and the data is available from GEO Accession GSE79257.^22^ The Novakovic study in was published 2019, and used the EPIC methylation array and the data is available from GEO under accession GSE131433^23^. Various assisted reproductive technology conception groups are described in these studies, but we only examined fresh in vitro fertilisation (IVF) conceived infants compared to naturally conceived infants. Raw intensity files (IDAT format) were read into R with minfi (v1.48.0),^24^ normalisation was conducted with the SWAN method,^25^ and probe filtering was conducted to remove probes with a detection p-value > 0.01 in addition to probes located on X or Y chromosomes. M values were computed and underwent differential methylation analysis using limma.^13^ The Estill study included 43 naturally conceived infants and 38 with fresh embryo transfer. The Novakovic study included 58 naturally conceived infants and 75 with fresh embryo transfer. Limma differential analysis was conducted accounting for sex, then methylation tables underwent LAM analysis with Reactome gene sets. Probe gene associations for HM450K array were established using the “IlluminaHumanMethylation450kanno.ilmn12.hg19” annotation set and the gene symbols were updated with HGNChelper. As the Novakovic study involved EPIC array data, gene-probe associations were obtained from the “IlluminaHumanMethylationEPICanno.ilm10b4.hg19” annotation set, and gene symbols were updated with HGNChelper.

### Differential pathway methylation in 19 common disease states

To examine whether LAM can identify differentially methylated pathways associated with disease prevalence and incidence, we downloaded blood-based EWAS summary statistics from a recent publication,^26^ made available via Zenodo (https://zenodo.org/records/8021411). The study used a longitudinal design, with sample collection baseline and diagnosis status established via electronic health record linkage up to 14 years after baseline. The study included 18,413 volunteers of European ancestry. We used summary statistics from the “full model”, which corrected for relevant potential confounders. Disease status at baseline was used to classify participants into prevalence groups. Disease status at follow-up was used to classify participants into incidence groups. The study used the EPIC Infinium array and 752,722 probes were described in the summary statistic dataset. As limma t-statistics were not available, we used probe delta-beta values as input for LAM. LAM was conducted individually for each disease condition with Reactome gene sets. Finally, as a test of type I errors, we randomised each of the 19 incident profiles using the base R sample command with a set seed, followed by LAM enrichment and counting the number of significant pathways. This process was repeated 100 times with unique seeds.

## Results

### Evaluation of FCS methods for Infinium methylation array data

Simulations were conducted using the eight FCS methods (LAT, LTT, LAW, LAM, LRM, ALT, ALW, ALM) and GSA, with group sizes of 3-12 and delta values between 0.1 and 0.5, and random gene sets with sizes 20, 50 and 100. At each parameter setting, the simulations were repeated 100 times with a different seed value. True positives, false positives and false negatives were used to calculate overall precision and recall at these three gene set sizes (**Figure 1A**). The LAM and ALM methods recorded an overall precision of 0.94, while GSA scored lowest with 0.848. Recall was highest for LAM with 0.22, while ALM scored 0.21 and GSA scored 0.15 F1 scores were highest for LAM with 0.36 followed by ALM with 0.35, while GSA scored 0.26.

**Figure 1.**
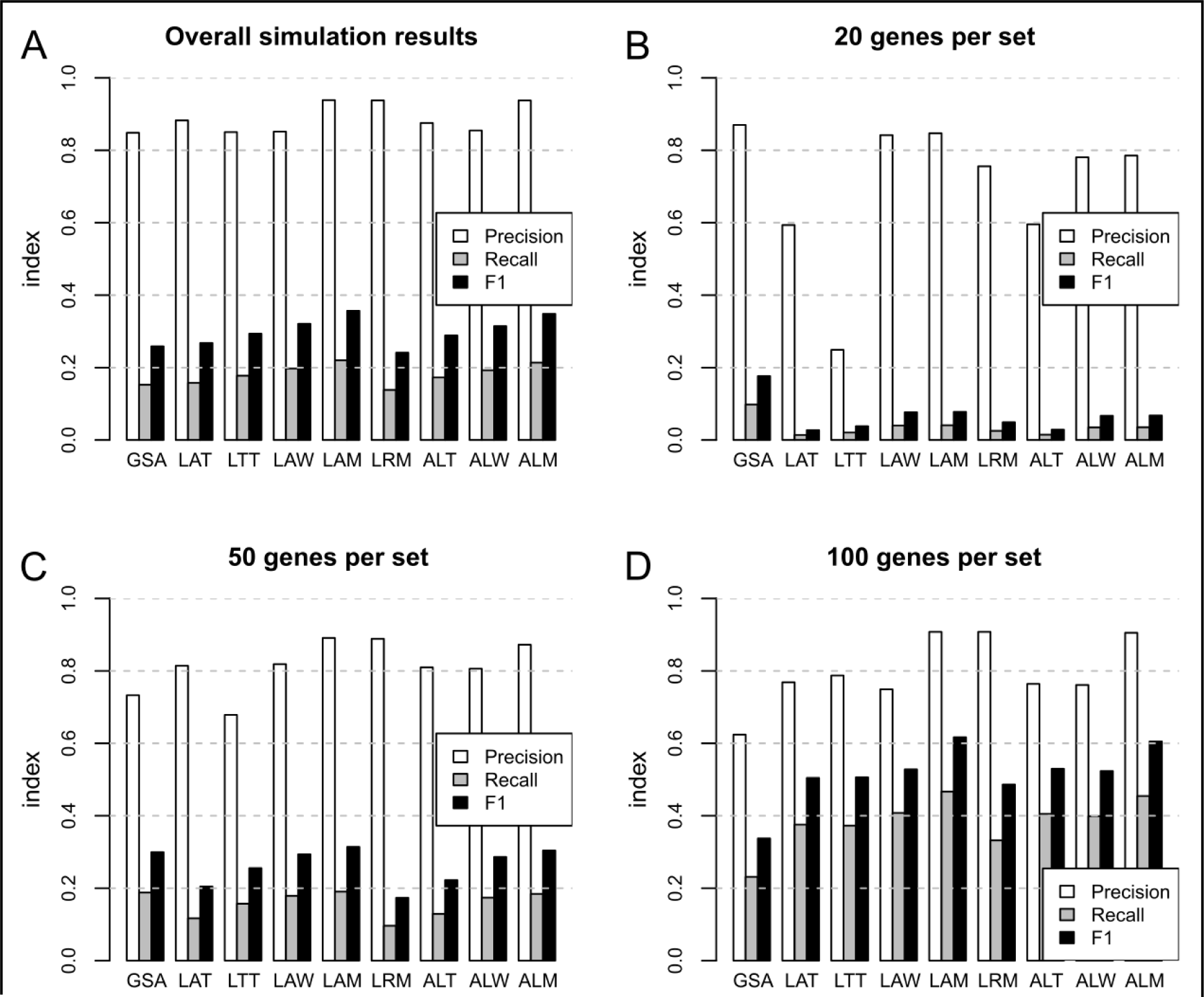
Precision, recall and F1 for enrichment tests with simulated data. (A) Overall results including 20, 50 and 100 genes per set. (B-D) Results for simulations using 20, 50 and 100 genes per set respectively.

We noticed that the size of gene sets used in the simulations strongly influenced the results, with overall recall increasing with gene set size (**Figure 1B-D**). The GSA method performed better than FCS methods with gene sets of 20 (**Figure 1B**), while at 50 genes per set GSA performance was on par with LAM **(Figure 1C**). At 100 genes per set, however, LAM showed superior precision and recall compared to GSA (**Figure 1D**).

Focusing on results from 100 genes per set, LAM precision was more consistent across the parameter ranges as compared to GSA which showed lower precision when group size and delta were lower (**Figure 2**). Recall was strongly dependent on group size and delta parameters. LAM recorded relatively higher recall when group size and delta were lower. F1 performance scores at 100 genes per set were better for LAM as compared to GSA.

**Figure 2.**
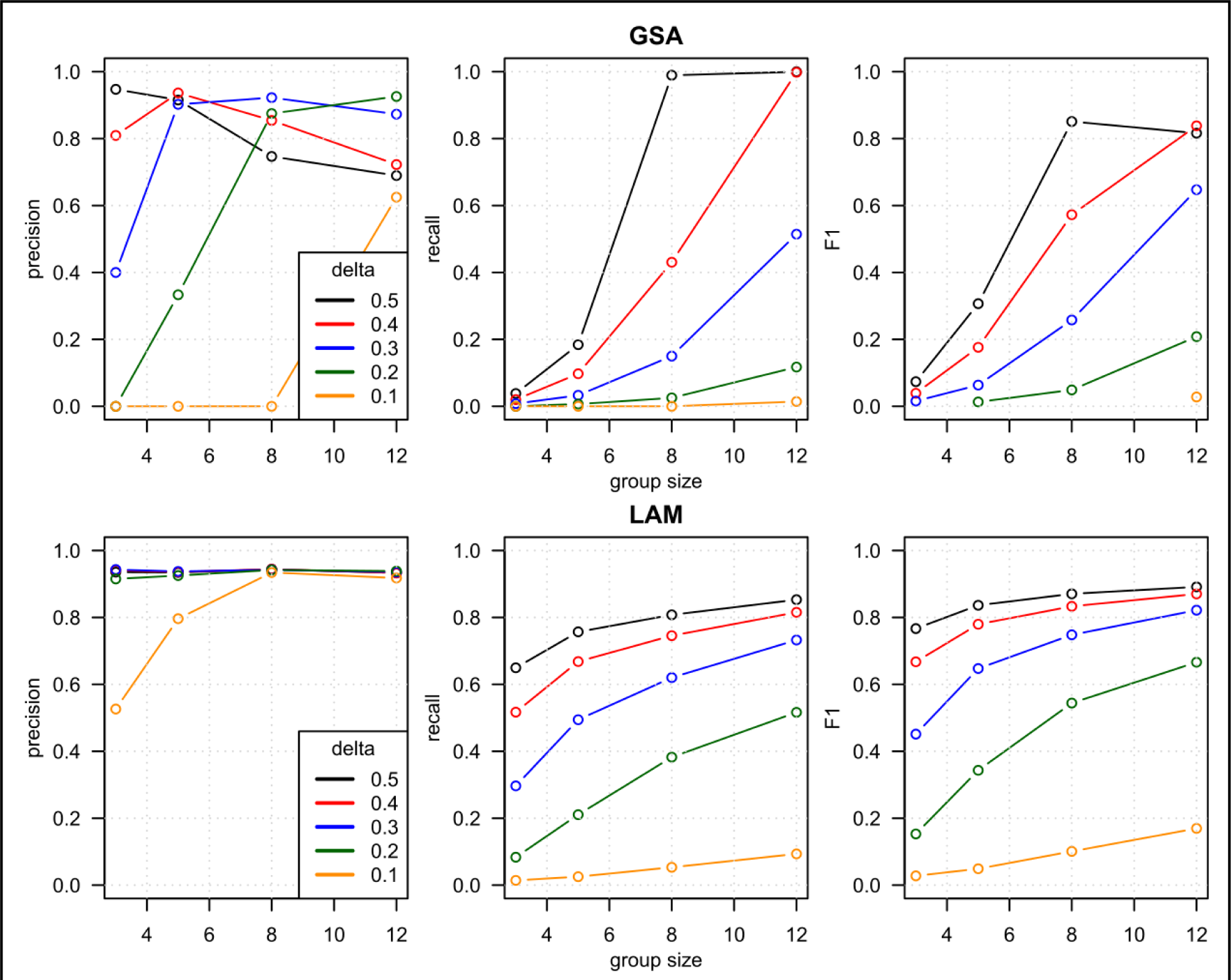
Precision, recall and F1 for GSA and LAM enrichment tests with simulated data over a range of group sizes and delta values. One hundred genes per set.

### Sensitivity of GSA and LAM methods with real cancer data

Next, GSA and LAM methods were applied to compare paired normal and cancer methylation data from 37 lung squamous cell carcinoma patients. From the 839,473 probes passing quality control, 473,572 were considered significant at 5% FDR. Of these, 119,785 probes were significantly increased in cancer, while 353,787 were significantly decreased. These significantly increased probes mapped to 16,520 unique genes, while the decreased probes mapped to 24,114 genes. These numbers are large, considering that there are 26,219 genes represented on this array. LAM resulted in 406 significant Reactome pathways at FDR<0.05. Of these, 370 involved higher methylation, while 36 involved lower methylation. LAM execution took 51 seconds using eight parallel threads. With GSA, there were 75 Reactome pathways with higher methylation and 32 with lower methylation. 12 pathways were significant in both directions. Thirty pathways were common between LAM and GSA methods. This includes 26 and 4 with higher and lower methylation respectively (**Figure 3A**). GSA execution for both directions in series took 22 seconds on one thread.

**Figure 3.**
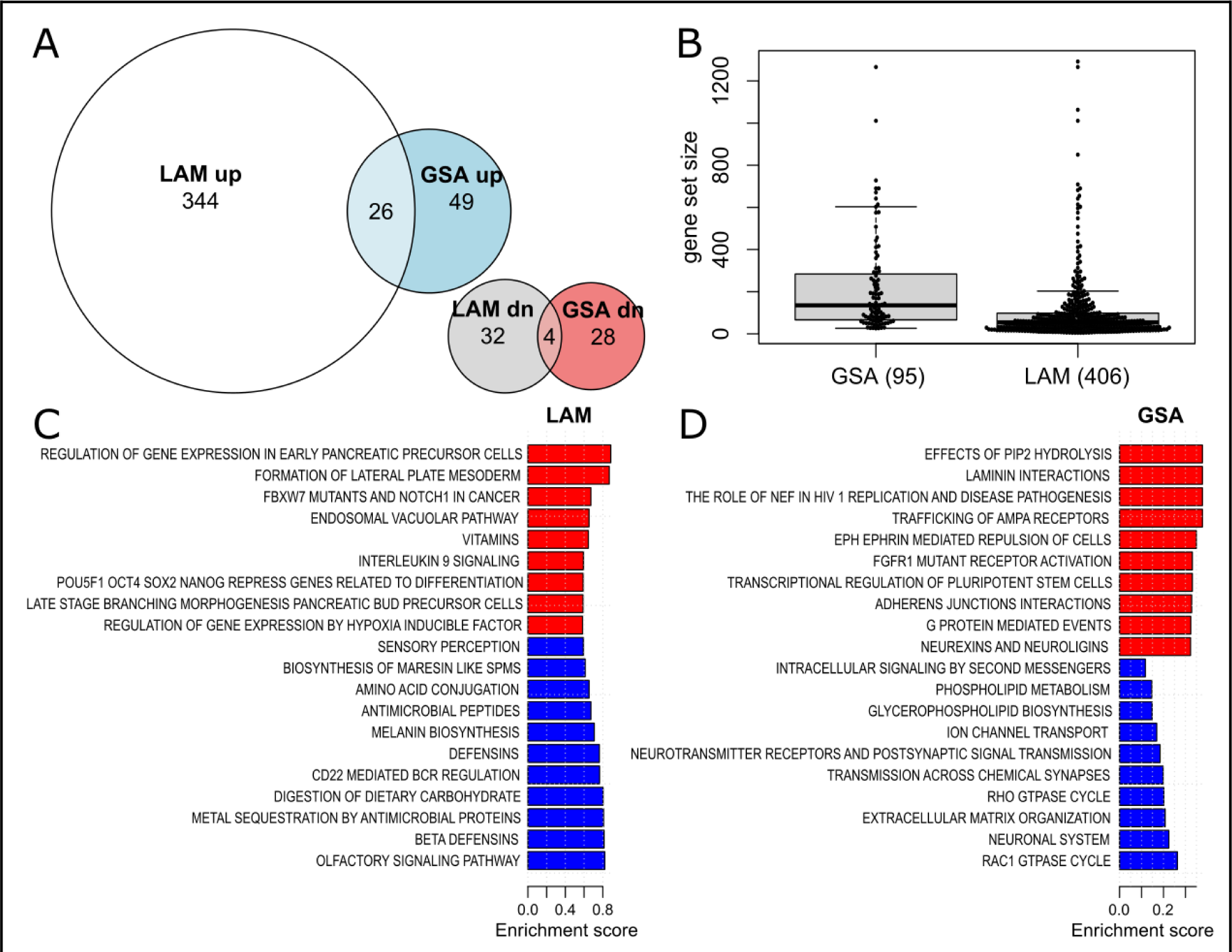
Pathway enrichment analysis comparing normal and cancer samples using LAM and GSA methods. (A) Euler diagrams showing the overlap in statistically significant pathways (FDR<0.05) from LAM and GSA results. (B) Sizes of significant gene sets found with LAM and GSA methods. (C) Significant LAM pathways with the largest S distance, an enrichment score generated by mitch and effect size proxy. (D) Top-ranked significant GSA pathways ranked by fold enrichment.

As the simulation results indicated LAM had better recall with larger gene sets, we were curious about whether there were differences in the sizes of gene sets found by LAM and GSA. GSA significant pathways had a median size of 135, while for LAM the median was 54.5, indicating pathways identified as significant with LAM were collectively smaller than GSA (**Figure 3B**).

Although algorithmic accuracy cannot be inferred from the types of pathways identified in real data,^27^ top-ranked results from LAM appeared to be more related to cell differentiation, identity and development (**Figure 3C**), as compared to GSA (**Figure 3D**).

To compare the sensitivity of these methods with real data, sample sizes were randomly downsampled, analysed as above and the significant pathways were compared to the results from the full group of 37 patients. This process was repeated 50 times at each sample size and the results are shown in (**Figure 4**). Pathways that were identified as statistically significant (FDR<0.05) in the smaller and full group were termed “consistent”, while pathways that were identified in the smaller but not in the full group were termed “inconsistent”. At a sample size of 10, LAM was able to detect 354 out of 406 (87%) consistent pathways, while GSA detected just 13 of 107 at this sample size (12%). LAM identified more inconsistent pathways than GSA, but the proportion of inconsistent findings was lower across all sample sizes. At a sample size of 10, the proportion of inconsistent pathways was 10% for LAM and 28% for GSA. This result suggests that LAM has superior sensitivity to detect differentially methylated pathways in real cancer data.

**Figure 4.**
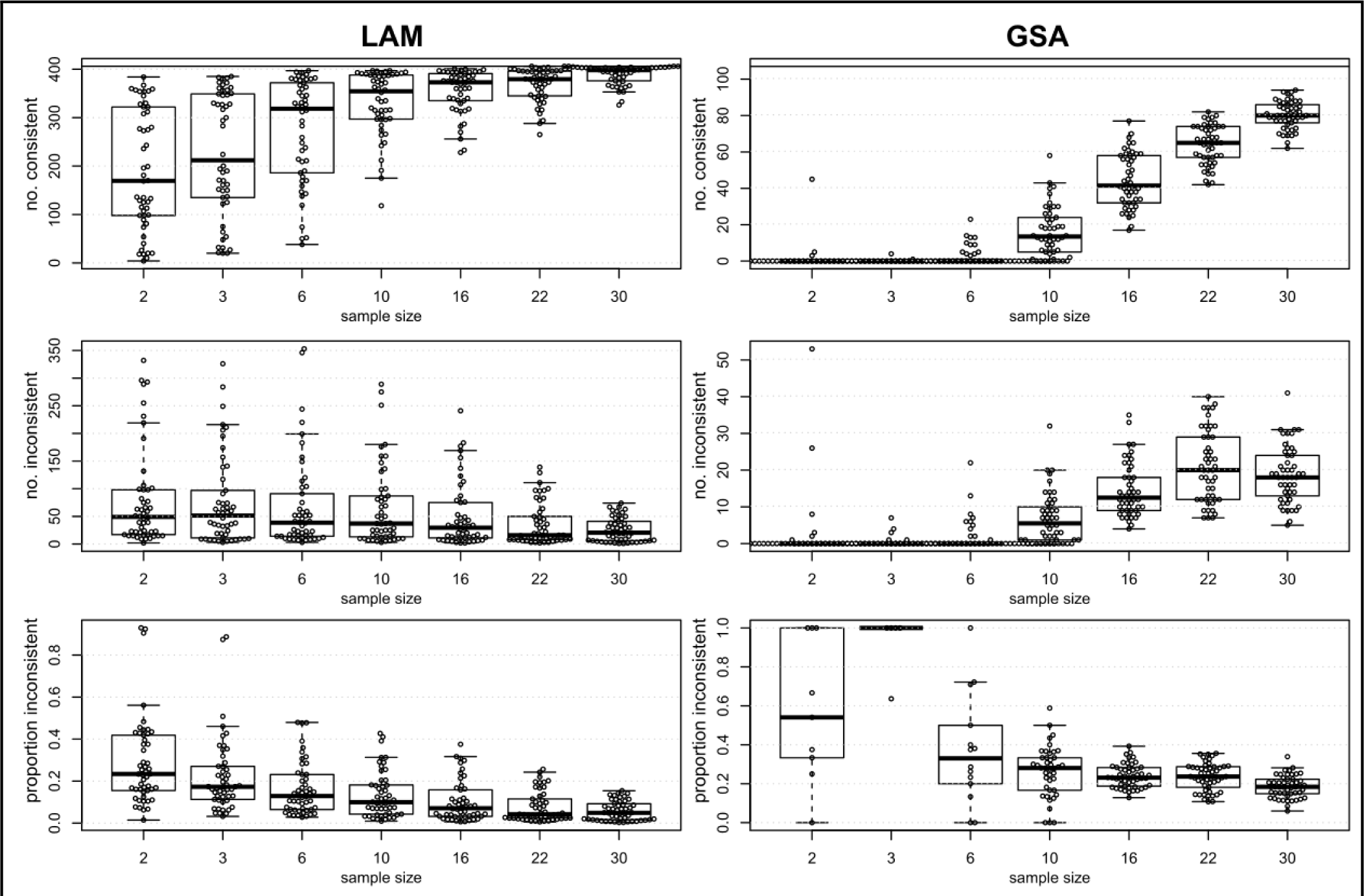
Sensitivity of LAM and GSA methods with real cancer array data. Upper panels show the number of significant pathways identified in the smaller sample that were consistent with the full group for LAM (left) and GSA (right). Middle panels show the number of significant pathways identified in the smaller sample that were inconsistent with the full group. Lower panels show the proportion of significant pathways identified that were inconsistent with the full group. Left panels correspond to LAM, and right panels to GSA methods.

### Integrating methylation and RNA expression pathways

Associations of epigenetic marks with gene expression are of great interest for understanding disease processes. Matching tumour-normal RNA-seq datasets were analysed and we conducted enrichment analysis of gene expression together with DNA methylation. From 18,704 detected genes, there were 12,380 differentially expressed genes (FDR<0.05), with 7,415 and 4,965 up-and down-regulated in the tumour group respectively. At the pathway level, there were 304 and 135 up-and down-regulated Reactome pathways with altered gene expression (FDR<0.05). As the context of gene methylation is important in influencing gene expression, promoter and gene body methylation were considered separately. In promoters, GSA identified 15 and 7 pathways with higher and lower methylation respectively, while at gene bodies, there were 18 and 75 pathways with higher and lower methylation respectively (FDR<0.05). Using LAM at promoters, there were 190 and 20 pathways identified with higher and lower methylation respectively. LAM at gene bodies identified 251 and 31 pathways with higher and lower methylation respectively (FDR<0.05). The relative overlap between significant promoter and gene body pathways was larger for LAM pathway sets (22%) as compared to GSA (3%) (**Figure 5A,B**). There was no observed overlap between GSA methylation-based pathways and gene expression pathways (**Figure 5A**), but there were some overlaps between pathways identified with LAM-based promoter and gene body methylation and gene expression (**Figure 5B**).

**Figure 5.**
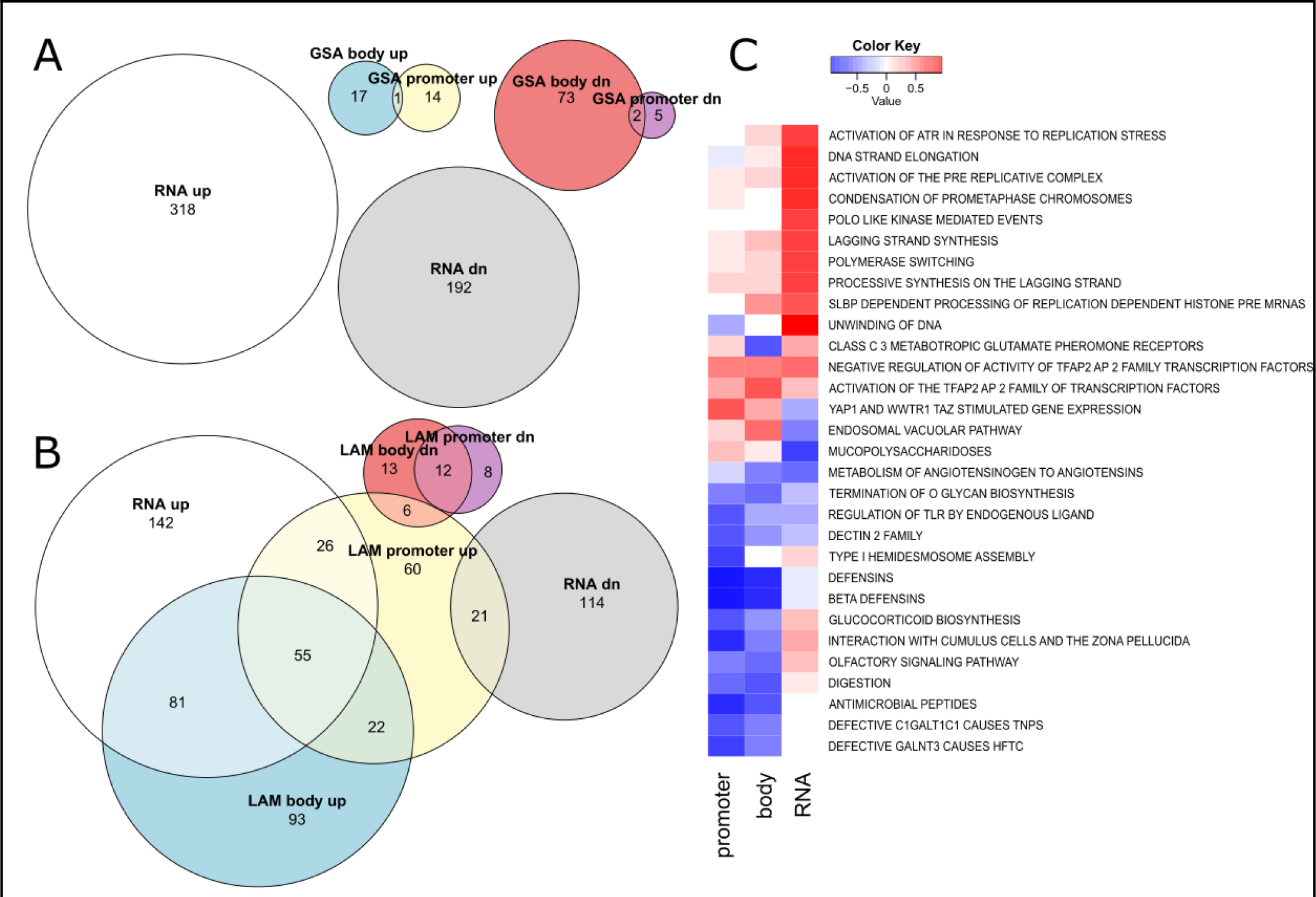
Integration of gene expression data with methylation pathways. (A,B) Euler diagrams showing the overlap in statistically significant pathways (FDR<0.05) from gene expression and GSA results (A) and LAM (B) results. (C) Multi-dimensional enrichment analysis of promoter and gene body methylation with gene expression using mitch. Top 20 gene sets shown with the largest S distance after FDR filtering at 0.05.

Using LAM, 136/304 upregulated RNA expression pathways were associated with increasing gene body methylation. Likewise, 21 pathways with downregulated RNA expression were associated with increasing promoter methylation. Some of these pathways with large enrichment scores across the three contrasts are depicted in heatmap form (**Figure 5C**). There were a small number of pathways that exhibited an inverse relationship between promoter methylation and RNA expression such as “unwinding of DNA”, “mucopolysaccharidoses”, “interaction with cumulus cells and the zona pellucida”, “glucocorticoid biosynthesis” and “olfactory signaling”. These were, however, less common than pathways that showed a positive relationship between RNA expression and promoter methylation. These results demonstrate that LAM can illuminate the complicated relationship between DNA methylation and gene expression at the pathway level in cancer.

### Exploration of methylation pathways in aging

We sought to explore the utility of LAM for exploring large EWAS datasets. One of the best-studied associations with DNA methylation is with chronological aging, although pathway-level differential methylation is not well defined. Using limma summary statistics from independent discovery and replication studies using the EPIC array platform described by McCartney et al (2020),^21^ we applied the LAM method to explore the pathway level differential methylation. In the discovery cohort, we observed 304 differential pathways (FDR<0.05), while in the replication group, we observed 107. There was a high degree of agreement, with 43 pathways significant in both groups. This is not surprising, as the gene-level rank differential methylation scores show a strong positive association between discovery and replication studies (**Figure 6A**). Using the rank-MANOVA test of the mitch package, we identified 390 pathways with altered methylation. Visualised as a scatterplot, the overall pattern is concordant, although a cluster of points in the lower right of the chart indicates 268 pathways with higher methylation in the discovery sample and lower methylation in the replication sample (**Figure 6B**). When prioritising results by S distance, the enrichment score reported by mitch and a surrogate measure of effect size, 29 of the top 30 pathways were concordant in their direction of regulation (**Figure 6C**). Some of the observed methylation changes make sense with reports in the literature. For example, the complement pathway is associated with aging^28^ and in this analysis we observe a reduction in methylation of genes belonging to this set including C1R, MBL2, C1S and C1QC (**Figure 6D**).

**Figure 6.**
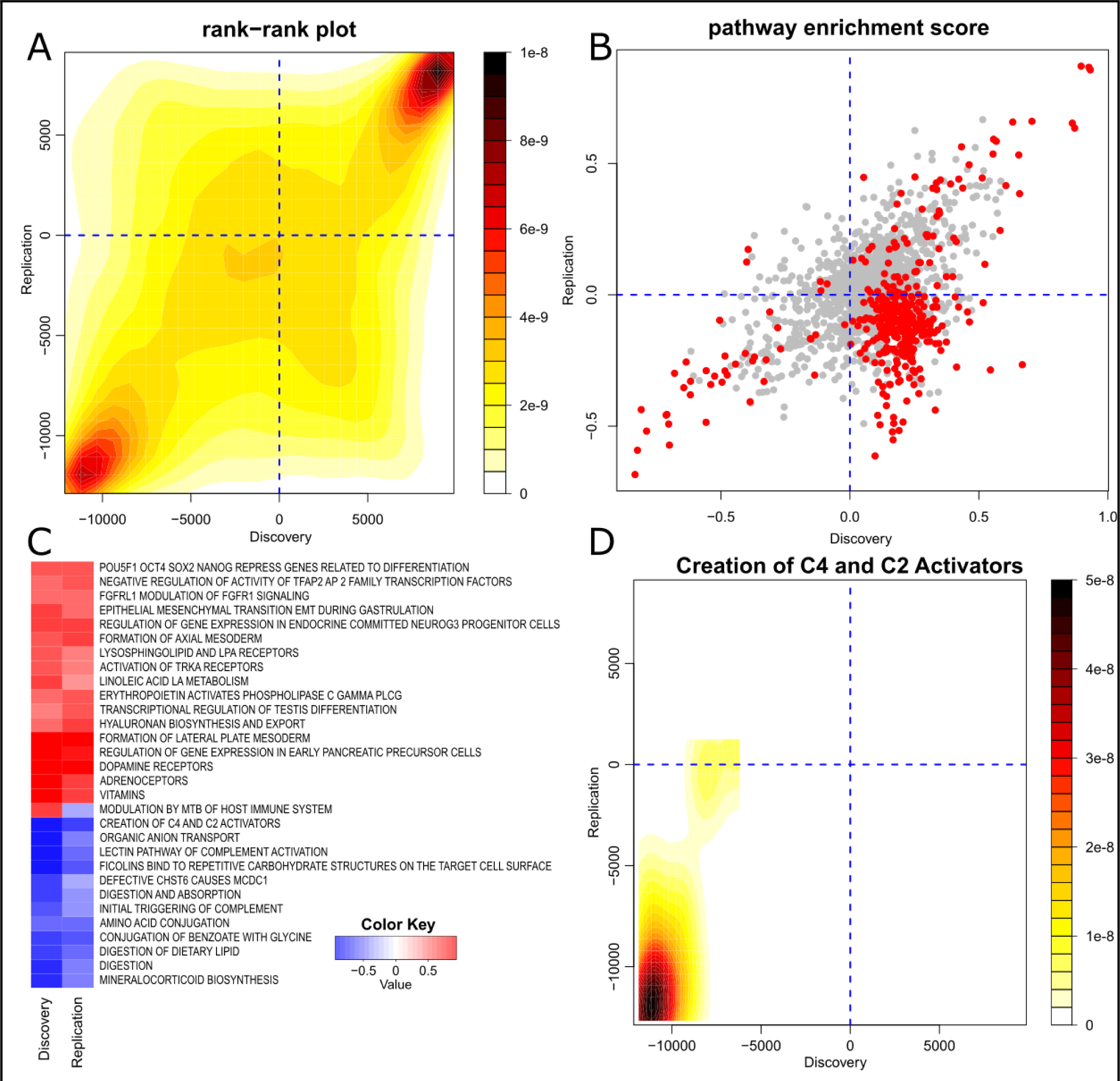
Pathway-level DNA methylation alterations with chronological age. (A) Contour heatmap showing the similarity in gene methylation score ranks in the discovery and replication studies. (B) Mitch pathway enrichment scores (S distances) in discovery and replication studies. Pathways with MANOVA FDR<0.05 are shown in red while others are shaded grey. (C) Heatmap of 30 pathways with most extreme S distances (a surrogate of effect size). Red indicates increasing methylation and blue shows lower methylation. (D) An example of a pathway identified with this method, “creation of C4 and C2 activators” shows lower methylation of member genes in both discovery and replication studies.

### Exploration of methylation pathway differences in infants conceived with in vitro fertilisation

Another application for this method is the joint pathway enrichment analysis of studies conducted with different array systems. Epigenetic differences between infants conceived naturally and with assisted reproductive technologies have been the subject of studies since 2009.^29^ Two of the highest-powered studies with publicly available data are Estill 2016 which used the HM450K array and Novakovic 2019 study which used the EPIC array.^22,23^ Joint enrichment analysis can uncover pathways that are common or different between these studies and illuminate the biological differences between conception groups. For this example, we focused on comparing naturally conceived infants to those conceived with in vitro fertilisation (IVF) with fresh embryo transfer. The Estill dataset on the HM450K platform had 418,833 probes that met the quality criteria (Natural n=43; IVF n=38). Of these, 149,589 probes showed statistically significant differential methylation (FDR<0.05), of which 28,767 and 120,822 exhibited higher or lower methylation in the IVF group respectively. The Novakovic dataset on EPIC array had 793,844 probes that met the filtering criteria (n=133). Of these, 5,562 were statistically significant (FDR<0.05) with 1,348 and 4,214 exhibiting higher or lower methylation in the IVF group respectively. After summarisation to the gene level, the Estill dataset on HM450K array describes 19,240 genes, while the Novakovic dataset on EPIC array describes 22,588 genes. There were 19,234 genes common to both platforms. The rank-rank plot of differential gene methylation scores indicates a high degree of similarity overall, and interestingly shows a trend of more genes having lower methylation levels in the IVF group (Figure 7A). A scatterplot of pathway enrichment scores indicates a moderate degree of agreement between these studies (*r*=0.35, *p*=2.2e-16) despite these studies being conducted on independent cohorts years apart and analysed with different array systems (Figure 7B). A heatmap shows a high degree of agreement of pathways with higher methylation in the IVF groups, while pathways with overall lower methylation in the IVF group showed some variability in enrichment scores between studies (Figure 7C). We were interested in which pathways had consistently lower methylation in the IVF group and identified the Reactome pathway “Adrenoreceptors” which had enrichment scores of -0.42 in the Estill study and -0.56 in the Novakovic study, recording a MANOVA FDR value of 0.04 (Figure 7D). Top-ranked genes from this pathway include *ADRA2C*, *ADRB2*, *ADRA2B* and *ADRA1B*. A loss of methylation of these genes could contribute to reported elevated blood pressure in people conceived by IVF.^30^ Taken together, this analysis shows that LAM method can be used to compare pathway enrichment across independent studies conducted with HM450K and EPIC arrays.

**Figure 7.**
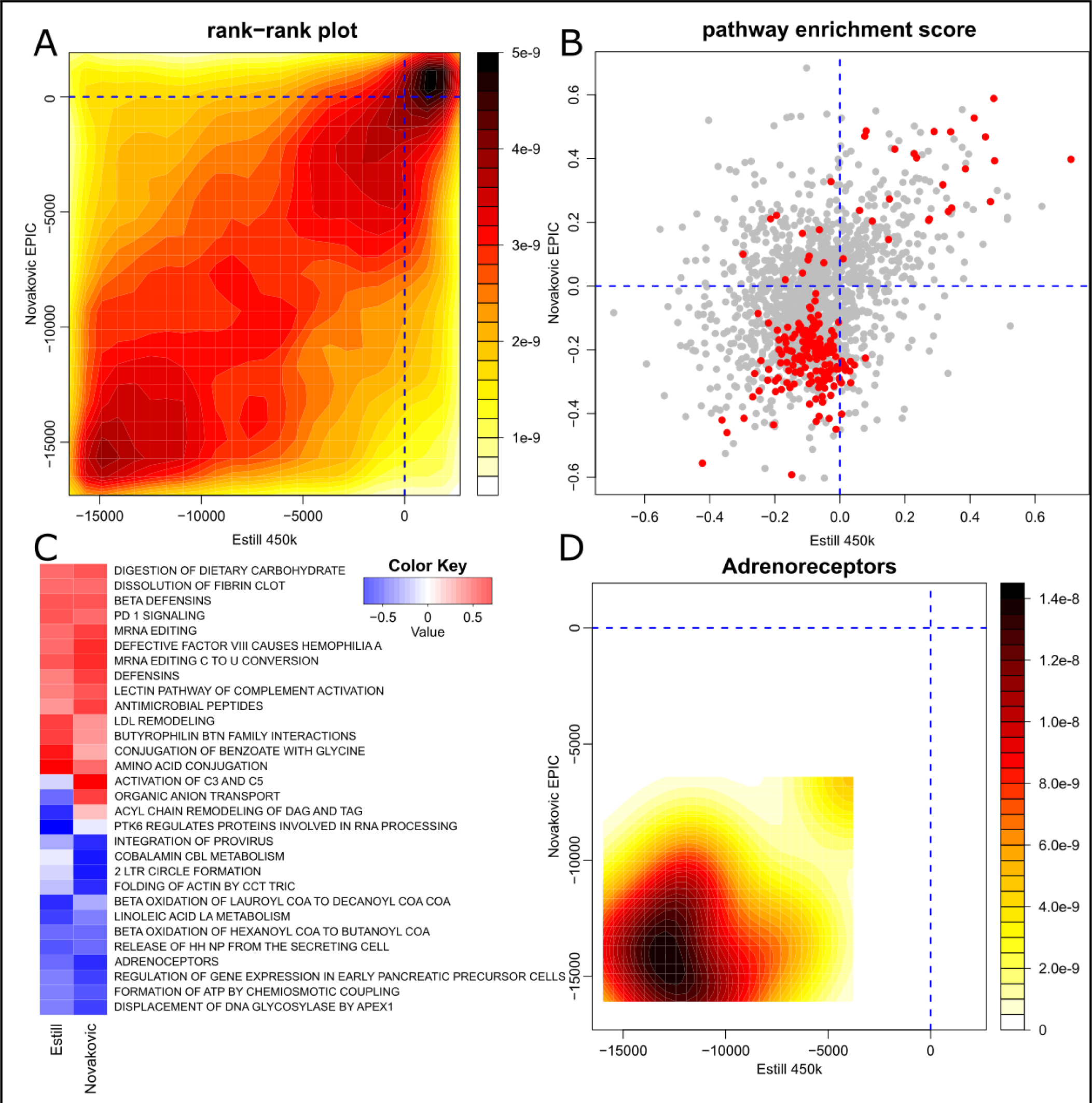
Pathway level DNA methylation differences in natural and IVF conceived infants. (A) Contour heatmap showing the similarity in gene methylation score ranks in the Estill (HM450K) and Novakovic (EPIC) studies. (B) Mitch pathway enrichment scores (S distances) in Estil and Novakovic studies. Pathways with MANOVA FDR<0.05 are shown in red while others are shaded grey. (C) Heatmap of 30 pathways with most extreme S distances (a surrogate of effect size). Red indicates higher methylation and blue shows lower methylation. (D) An example of a pathway identified with this joint enrichment analysis method, “Adrenoreceptors” shows lower methylation of member genes in both Estill and Novakovic studies.

### Differential pathway methylation in 19 common disease states

To examine the potential for LAM to reveal differential pathway methylation associations with diseases, we obtained and analysed blood-based EWAS summary statistics from a recent publication.^26^ This study examined methylation in 14 prevalent disease states and the incidence of 19 disease states in a group of 18,413 participants, using the EPIC array. Delta-beta values were used for the purpose of scoring probe differential methylation, and these values underwent LAM analysis with Reactome gene sets.

There were 899 differentially methylated pathways in the prevalence arm of the study (FDR<0.05) (Figure 8A). On average, there were 64 pathways with differential methylation in each prevalent condition, with Chronic obstructive pulmonary disease (COPD) having the most (263) and Alzheimer’s disease having the fewest (3). We selected up to five statistically significant pathways with an absolute S distance of >0.4 in either direction in each prevalent condition to include in a heatmap (Figure 8B). We observed reduced methylation in the angiotensinogen metabolism pathway in patients with diabetes. Patients with colorectal cancer had lower methylation of the interleukin 10 signaling pathway. Reduced methylation of the salty taste perception pathway was observed in patients with CKD.

**Figure 8.**
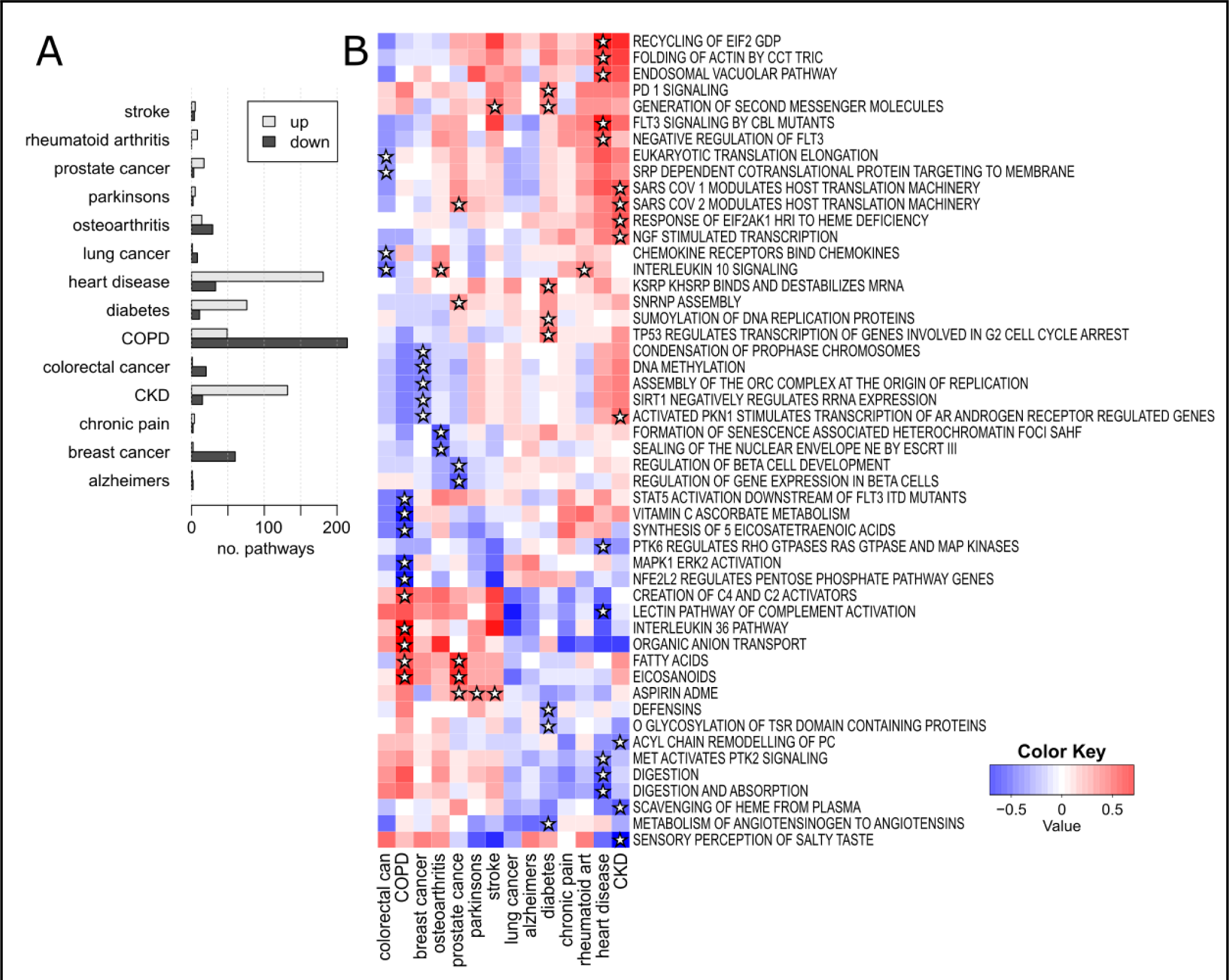
Differential pathway methylation associated with prevalence of 14 common disease states. (A) A bar plot showing the number of statistically significant pathways with higher and lower methylation identified in each prevalent condition (FDR<0.05). (B) A heatmap of S scores for selected pathways across 14 common prevalent disease states. Stars indicate that the pathway was identified as being among the top five differentially methylated pathways in each direction for each condition.

We also wanted to know whether the LAM method could identify signatures that appear before patients are diagnosed with a condition. Therefore, we examined differential pathway methylation associated with the incidence of 19 disease states. Across all 19 conditions, there were 1570 significant pathways (FDR<0.05) (Figure 9A). Liver cirrhosis had the most, with 464, followed by COPD with 400, while Alzheimer’s had just one and COVID-19 hospitalisation had none. A heatmap shows the enrichment scores for selected pathways across these incident conditions (Figure 9B). Breast cancer incidence was associated with reduced methylation to the metal sequestration pathway. Incident prostate cancer was associated with reduced methylation of the FGFR1 pathway. Incident Parkinson’s disease was associated with reduced methylation of aquaporins. In COPD incidence, higher methylation of carbohydrate-binding ficolins was observed together with reduced methylation of the GDP mannose synthesis pathway. Incident stroke was associated with higher methylation of the lipoxin synthesis pathway.

**Figure 9.**
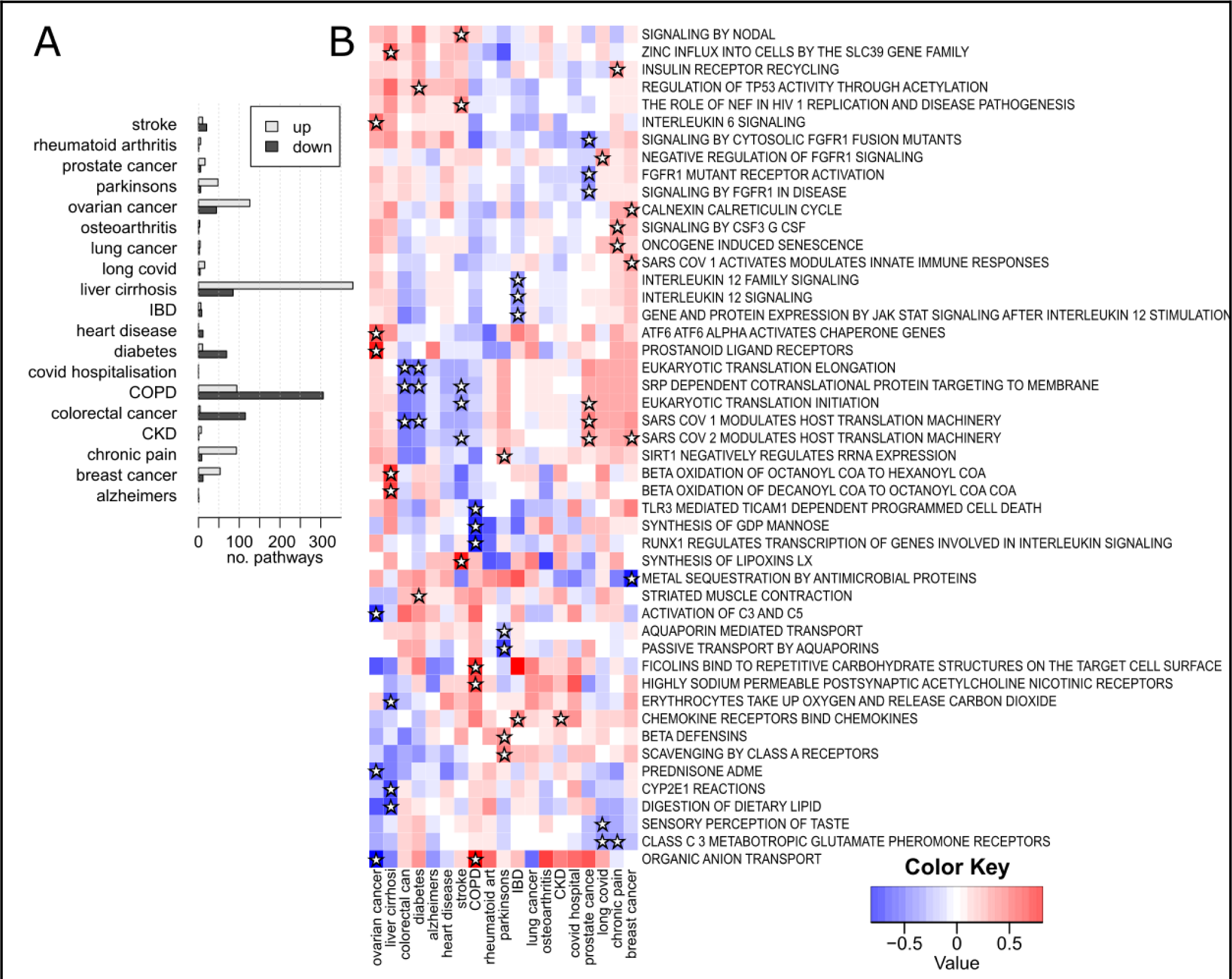
Differential pathway methylation associated with incidence of 19 common disease states. (A) A bar plot showing the number of statistically significant pathways with higher and lower methylation identified in each incident condition (FDR<0.05). (B) A heatmap of S scores for selected pathways across 19 common incident disease states. Stars indicate that the pathway was identified as being among the top three differentially methylated pathways in each direction for each condition.

To show that these findings are not the result of false positives from the LAM method, we randomised the incidence profiles for all 19 conditions prior to enrichment analysis. This was repeated 100 times, and these analyses of randomised data yielded few significant pathways. Specifically, from 100 runs 64 yielded no significant pathways across 19 conditions. Twenty-eight repeats had fewer than five false positives and eight runs had more than five false positives. Across the 100 repeats, the mean number of false positives was 1.95 and the median was 0. These results indicate that LAM can identify differentially methylated pathways in disease groups and detect altered methylation before a diagnosis has been made, with few false positives.

## Discussion

Overall, we found that the FCS methods performed better across most simulation conditions. Of the different FCS methods tested, LAM was superior in terms of precision and recall. LAM also fits relatively easily into existing workflows, as most EWAS studies seem to use limma in their analyses of differential probe methylation. The ALM method had only slightly worse performance as compared to LAM, but ALM requires users to aggregate the probe methylation values for each sample for each gene before limma. This extra aggregation step is computationally intensive and not a standard part of existing methylation analysis workflows.

Our analyses also consistently showed that non-parametric methods showed better recall with only a small decrease in precision. Summarisation of probe level t-statistics to gene level only takes a few seconds with the efficient base R aggregate command, and once aggregated, signatures can be analysed with existing packages like mitch. Mitch does not require a large amount of system memory and due to its parallel architecture can make use of multi-threaded processors, so it is not much slower than conducting ORA with existing methods that are single-threaded.

An interesting observation is that ORA performed relatively better when gene sets were smaller in our simulations. Given that popular pathway databases like KEGG and Reactome pathways consist of both large (>100) and small (<20) gene sets, a hybrid FCS and ORA approach might increase sensitivity and recall.

At 100 genes per set, LAM had superior sensitivity compared to GSA, which agrees with a previous report of mitch’s performance with simulated RNA-seq gene expression data^14^. The improved sensitivity of the LAM method provides a new opportunity for researchers to reanalyse previously conducted EWASs with contemporary pathway databases to better understand subtle signatures, as we have demonstrated with the chronological age, assisted reproduction examples and 19 disease EWAS analyses. The analyses conducted here used Reactome pathways, but LAM is general-purpose, so different gene sets can be used to examine various hypotheses. Transcription factor target gene sets can be used to identify transcription factors associated with changes in pathway methylation. MicroRNA target gene sets can identify potential associations between microRNA targets and DNA methylation patterns. The MSigDB resource contains these and several other types of gene sets for exploration.^30^

The differences in pathway results obtained with LAM and GSA from lung cancer data were striking, not only in the number of pathways identified, but also in their relevance to the disease. Interestingly, the relationship between promoter methylation and gene expression was not always inverse as we had anticipated. Joint enrichment analysis showed some pathways with increasing promoter methylation together with increasing RNA expression (e.g., “Negative regulation of AP2 transcription factors”). Likewise, a few pathways showed decreases in methylation with decreases in RNA expression such as “Dectin family”, “TLR regulation” and “glycan biosynthesis termination”. These results are in line with previous reports of the widespread existence of both negative and positive relationships between DNA methylation and gene expression^31–34^, and support the idea that there is a complicated relationship between DNA methylation and gene expression in lung cancer.

We have demonstrated that this method works with both HM450K and EPIC array data and can even facilitate the comparison of EWAS across different platforms. We have provided an example workflow that researchers can use as a template. It shows how to generate a probe-gene table from Bioconductor annotation datasets, how to update the defunct gene names and how to run the enrichment analysis. Using this example, generating similar gene tables for the new Infinium Mouse and EPIC 2.0 arrays and any future arrays will be straightforward, once their annotation sets are available through Bioconductor.

Updating defunct gene symbols is an important step as 2,788 and 3,253 gene symbols were updated on the HM450K and EPIC array annotations, which represent approximately 10-14% of genes on these platforms. We note that other packages for enrichment of Infinium array data do not by default address this, which could lead to a loss of these genes from pathway enrichment and diminished sensitivity.

The analysis of the large-scale disease EWAS shows that LAM can be readily applied to identify differential pathway methylation associated with prevalent and incident disease. The authors of the study used GOmeth for their pathway analysis and from the 33 models examined, significant findings were obtained for only two, prevalent diabetes and heart disease. In contrast, LAM identified 2,469 pathway associations (74.8 per model). To demonstrate these are not the results of false positives, permuted profiles yielded on average 0.10 significant findings per model, supporting the idea that LAM is identifying real pathway signatures.

This work could be extended to include not just proximal CpG sites, but also enhancer-based probes and their target genes based on resources such as GeneHancer.^35^

There is also potential to generate new libraries of differentially methylated probes and gene sets, to contribute to the pool of molecular signatures in public resources like MSigDB, which will assist in understanding similarities between methylation profiles.

## Data availability

Data sets reanalysed here are publicly available from NCBI GEO, Zenodo or from the supplementary tables of journal articles as described in the Methods section.

## Code availability

Code to generate the results shown here are included in a GitHub repository (https://github.com/markziemann/gmea/). To enable reproduction of the findings, a docker image is made available at DockerHub (https://hub.docker.com/r/mziemann/gmea). A reproducible example workflow report is hosted on our website (https://ziemann-lab.net/public/gmea/example_workflow.html). These materials have been archived at Zenodo (https://doi.org/10.5281/zenodo.10685538).

## Acknowledgements

We thank the researchers who publicly shared data to enable this work.

